# Motion Coherence and Luminance Contrast Interact in Driving Visual Gamma-Band Activity

**DOI:** 10.1101/741066

**Authors:** Franziska Pellegrini, David J Hawellek, Anna-Antonia Pape, Joerg F Hipp, Markus Siegel

**Affiliations:** Hertie Insitute for Clinical Brain Research, University Tübingen, Tübingen, Germany; Center for Integrative Neuroscience, University Tübingen, Tübingen, Germany; MEG Center, University Tübingen, Tübingen, Germany; Roche Pharma Research and Early Development, Neuroscience and Rare Diseases, Roche Innovation Center Basel, Basel, Switzerland

## Abstract

Synchronized neuronal population activity in the gamma-frequency range (> 30 Hz) correlates with the bottom-up drive of various visual features. It has been hypothesized that gamma-band synchronization enhances the gain of neuronal representations, yet evidence remains sparse. We tested a critical prediction of the gain hypothesis, which is that features that drive synchronized gamma-band activity interact super-linearly. To test this prediction, we employed whole-head magnetencephalography (MEG) in human subjects and investigated if the strength of visual motion (motion coherence) and luminance contrast interact in driving gamma-band activity in visual cortex. We found that gamma-band activity (64 to 128 Hz) monotonically increased with coherence and contrast while lower frequency activity (8 to 32 Hz) decreased with both features. Furthermore, as predicted for a gain mechanism, we found a multiplicative interaction between motion coherence and contrast in their joint drive of gamma-band activity. The lower frequency activity did not show such an interaction. Our findings provide evidence, that gamma-band activity acts as a cortical gain mechanism that nonlinearly combines the bottom-up drive of different visual features in support of visually guided behavior.

## Introduction

Synchronized neuronal population activity in the gamma-frequency range (> 30 Hz), i.e. gamma-band activity, is a hallmark of feed-forward visual processing (Donner and Siegel 2011; Vinck et al. 2013; van Kerkoerle et al. 2014; Fries 2015). It is robustly driven by sensory stimulation and varies with several parameters of visual stimuli such as stimulus size (Gieselmann and Thiele 2008; Perry et al. 2013; Vinck and Bosman 2016), luminance contrast (Hall et al. 2005; Henrie and Shapley 2005; Niessing 2005; Ray and Maunsell 2010b; Hadjipapas et al. 2015; Perry et al. 2015), stimulus orientation (Friedman-Hill 2000; Siegel and König 2003; Koelewijn et al. 2011) and visual motion (Liu and Newsome 2006; Siegel et al. 2007; Muthukumaraswamy and Singh 2013). Gamma-band activity increases monotonically with visual motion coherence (Siegel et al. 2007) and increases approximately linearly with luminance contrast (Hall et al. 2005; Henrie and Shapley 2005; Niessing 2005; Ray and Maunsell 2010b; Hadjipapas et al. 2015; Perry et al. 2015).

Gamma-band activity is also related to cognitive processes. It correlates with selective visual attention (Fries 2001; Siegel et al. 2008), and predicts visual discrimination performance (Siegel et al. 2008) and reaction times during sensory discrimination (Womelsdorf et al. 2006; Rohenkohl et al. 2018). Thus, gamma-band activity may reflect a cortical gain mechanism. Enhanced rhythmic synchronization may increase the impact of neuronal spiking onto downstream neuronal populations in the context of visually guided behavior (König et al. 1996; Salinas and Sejnowski 2001; Fries et al. 2007; Donner and Siegel 2011; Fries 2015).

A critical prediction of the cortical gain hypothesis is that a combination of visual features that drive gamma-band activity does result in a super-linear (e.g. multiplicative) interaction, rather than a mere additive effect of these features on gamma-band activity. We tested this prediction recording magnetencephalography (MEG) in human participants that viewed dynamic random-dot motion patterns with varying luminance contrast and motion coherence. We found that, in addition to a linear drive of gamma activity through coherence and contrast, these stimulus features indeed showed a multiplicative interaction. Modulations of gamma-band activity were localized to visual cortex and accompanied by a more widespread modulation of lower frequency activity (8 – 32 Hz) that did not show an interaction between stimulus features. Our results provide novel evidence that gamma-band activity reflects a bottom-up driven cortical gain mechanism in the support of visually guided behavior.

## Material and Methods

### Participants

Nineteen subjects (5 male, mean +− SD age, 26.2 +− 3.2 years; age range, 21-35 years) participated in the experiment and received monetary compensation for their participation. The study was conducted in accordance with the Declaration of Helsinki, approved by the local ethics committee and informed consent was obtained from all subjects prior to the recordings. All subjects were in good health and had normal or corrected-to-normal vision. 14 subjects participated in the full experiment of 900 trials, one subject stopped after 283 trials, one subject stopped after 798 trials, one subject after 881 trials, one subject after 894 trials and one subject after 899 trials.

### Stimulus material

The stimuli consisted of dynamic random-dot patterns with bright dots on a black background (Fig. 1). During the entire experiment, subjects sat in the MEG in upright position. For every trial, they first saw a blank black screen with a fixation cross in the center (Fig. 1A). After 500 ms, a dynamic random-dot stimulus appeared on either the left or the right side of the fixation cross (10 deg eccentricity, 12 deg stimulus diameter). 1000 ms later, the stimulus disappeared. After a variable delay (300 to 600 ms) a Go cue was given through a brief dimming of the fixation cross (1 frame, ~16 ms). The participants then pressed one of two buttons to indicate whether they saw an upward or downward stimulus motion.

**Figure 1.**
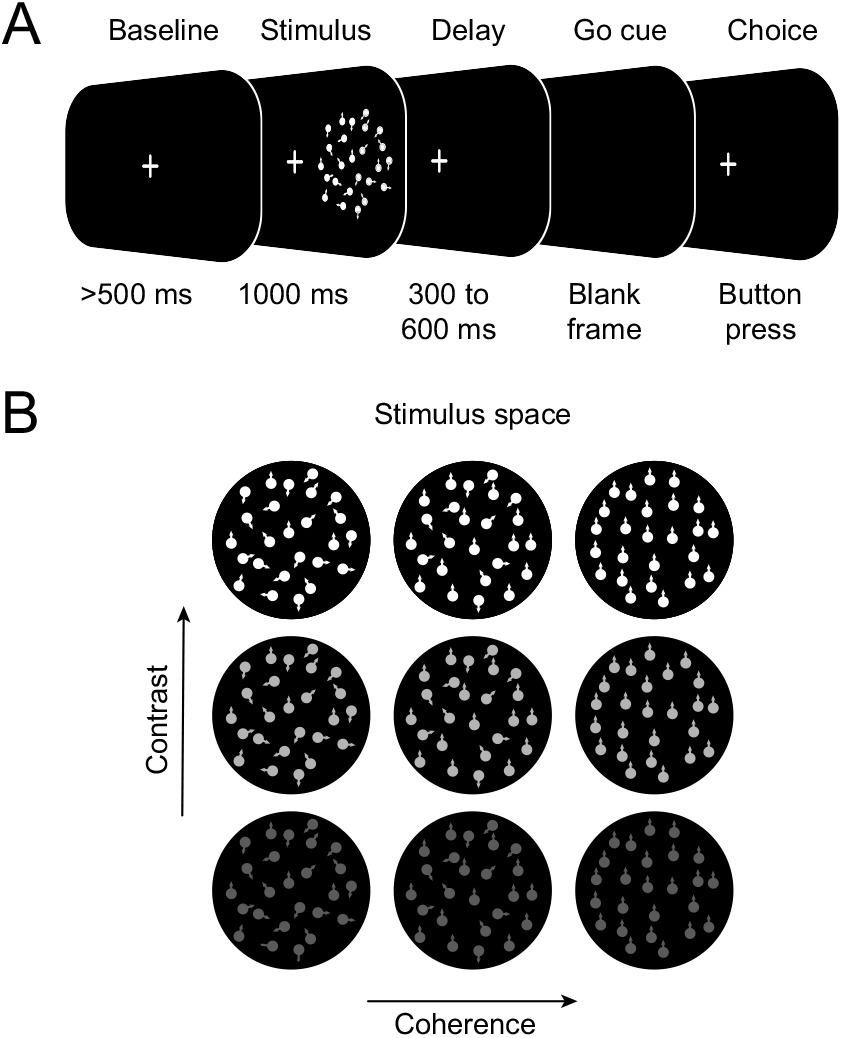
Experimental paradigm and stimulus space. *A*, On each trial, subjects fixated a small central fixation cross. Following a blank baseline (>500 ms), a dynamic random-dot stimulus appeared either on left or right side and disappeared again after 1000 ms. After a variable delay (300-600 ms), the fixation cross disappeared as a go cue for the response. Subject reported the perceived motion direction (up vs. down) with a button press (left vs. right hand) *B*, Stimuli varied in motion coherence (12%, 56%, 100%) and luminance contrast (20%, 60%, 100%).

The stimuli had varying features with three levels of motion coherence (12%, 56%, 100%; Fig. 1B) and three levels of luminance contrast (Weber contrast levels: 20%, 60%, 100%). The varying coherence was induced using random direction noise with the ‘same’ rule (Scase et al. 1996) (12 deg stimulus diameter, 1000 dots, 10 deg/s dot speed, 0.1 deg dot radius). Thus, a fraction of dots — the ‘signal dots’ — moved all in the signal directions (upward or downward movement) during the entire stimulus presentation, while another fraction of ‘noise dots’ moved in random directions. We manipulated the contrast of the stimuli by increasing dot brightness against a constant luminance background. The task factors motion direction (upward vs. downward), presentation side (left vs. right), coherence level and contrast level were counter-balanced and randomly varied across trials during every experiment. The stimulus-response mapping (button press with the left vs. right hand to indicate upward vs. downward motion, respectively) was counter-balanced across subjects.

### Data acquisition and preprocessing

MEG was continuously recorded using a 275-channel whole-head system (Omega 2000, CTF Systems Inc.) in a magnetically shielded room. The head position relative to the sensors was measured using three head localization coils (nasion, left/right pre-auricular points). Electroencephalography (EEG) recordings were performed in parallel to the MEG recordings. The EEG data is not presented here. All analyses were performed in Matlab (MathWorks) using custom code and the open source toolboxes Fieldtrip (Oostenveld et al. 2011) and SPM12 (http://www.fil.ion.ucl.ac.uk/spm). The MEG signals were recorded with a sampling rate of 2483.8 Hz. Off-line, the data was high-pass filtered with a cut-off frequency of 1 Hz and down-sampled to 500 Hz. Line noise was removed by applying band-stop filters at 50, 100, 150, 200 and 250 Hz with cut-offs at 1 Hz (all 4th-order zero-phase Butterworth filters). Trials containing jumps and channels that were affected from flux trapping due to the simultaneous EEG-MEG recordings were excluded from the analysis. We conducted an independent component analysis (FAST ICA; Hyvärinen and Oja 2000) to further clean the data from eye blink, eye movement, muscular and pulse artefacts. We inspected the first 100 components of each subject visually according to their topology, time courses and spectra. Components that could be clearly detected as artefacts were subtracted from the data before further analysis (mean: 4.7; SD: 4.1 components per subject).

### Spectral Analyses

All spectral analyses were performed using Morlet’s wavelets. We retrieved the time–frequency representation (TFR) for frequency f of the signal x(t) at time t with the convolution operation:

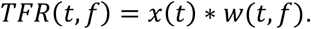

We write w(t,f) for Morlet’s wavelets:

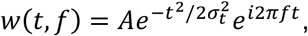

where *σ_t_* is the standard deviation (*SD*) of the signal in the time domain and *A* is a normalization factor. In both time and frequency domain, Morlet’s wavelets are Gaussian shaped. *σ_f_* = *q/f* is the *SD* in the frequency domain at frequency *f* and *q* is the width of the wavelet. The *SD* in time domain is given by *σ_t_* = 1/(2*πσ_f_*). We used a spectral width of 5 cycles and a temporal width of 3*σ_t_*. We computed the wavelet transformation in steps of 20 ms at frequencies between 8 and 256 Hz, logarithmically scaled in quarter octave steps.

### Source localization

We projected the frequency-decomposed MEG data to predefined source locations using adaptive linear spatial filters (Beamforming) (Van Veen et al. 1997; Gross et al. 2001). To account for different head anatomy, we constructed a personalized lead field for each participant individually. The lead field describes what signal we would observe at the sensors, if an isolated dipole with fixed current pointing to each of the 3 principle axes was active (forward model). The construction of lead fields was based on T1 magnetic resonance imaging (MRI) data from each subject. First, we segmented the data into different tissue types: grey and white matter, cerebrospinal fluid, skull and skin. Based on the segmented MRI data, we constructed individual single-shell head models (Nolte 2003) and subsequently matched a standardized source-model to the individual brain shapes (Hipp et al. 2012). The source-model contained 400 locations that homogeneously covered the space below the MEG sensors approximately 1 cm beneath the skull. Source coordinates, head model and MEG channels were calculated relative to the three head localization coils. We used DICS beamforming in the frequency domain (Gross et al. 2001) to project the data from sensor level to source space. Beamforming renders activity from sources of interest with unit gain, while suppressing contribution from all other sources. Briefly, DICS beamforming uses the cross-spectral density matrix (*CSD*) on sensor level, specifically for every frequency band, and the individual lead fields *L* to define spatial filters *F*:

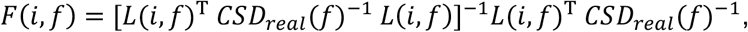

where i denotes sources and *CSD_real_* is the real part of the *CSD*. To obtain the 3-by-3 source level *CSD* (*CSD_i_*), we projected the sensor level CSD through the filter *F*:

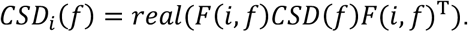

We then performed principal component analysis (singular value decomposition) on the source level *CSD_i_* and selected the first principal component that represents the most dominant dipole orientation. Subsequently, we projected the Filter *F* onto the first principal component and obtained *F_pri_*. Finally, we projected the *TFR* data from the sensor level to source level by multiplying them with the filter *F_pri_*:

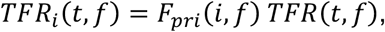

where *TFR_i_* denotes the time–frequency representation on source-level.

### Response normalization

To suppress stimulus-evoked responses, we subtracted the average potential across trials per condition, subject, time-frequency bin and source.

To project the data into a consistent relationship between cortical hemisphere and side of stimulation (left vs. right), we flipped the responses across the sagittal axis of the brain. Thus, the left hemisphere of the brain represents activity contralateral to the stimulation and the right hemisphere represents the activity ipsilateral to the stimulation.

We characterized spectral responses *R*(*f*) as the percentage of change in signal amplitude at frequency f and time t relative to the pre-stimulus baseline:

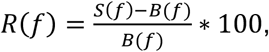

where *S*(*f*) denotes the spectral amplitude in the temporal interval of interest and *B*(*f*) denotes the spectral amplitude during the pre-stimulus baseline (500 ms before up to stimulus onset), averaged across all trials and conditions. We show the spatial distribution of power relative to baseline in five frequency bands, averaged across all subjects, trials and time bins from 0.1 to 1.1 s post stimulus onset. At later time points, we observed activity modulations that localized to the motor cortex, which likely reflected neuronal process related of response preparation. Thus, we restricted all analyses to the time window from 0.1 to 0.6 s post stimulus onset. In order to assess the modulation of neuronal responses by stimulus conditions (see below), we normalized all neuronal activity according to the mean across all trials and conditions of a subject, separately for every time, frequency and source bin.

### Analysis of Response Modulation

We used sequential polynomial regression (Büchel et al. 1998; Rees et al. 2000; Siegel et al. 2007) to assess the modulation of responses by contrast and coherence. The response *y* was modelled as a combination of predictor variables *x*_1_ (contrast) and *x*_2_ (motion coherence):

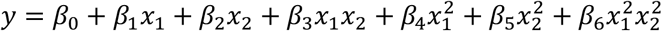

with *β* reflecting the polynomial coefficients. To assess the amount of variance that each predictor accounted for independently, we orthogonalized the regressors prior to model fitting using QR-decomposition. Starting with the zero-order (constant) model and based on F-statistics (Draper and Smith 1998), we tested whether incrementally adding new predictors improves the model significantly. We used the identical procedure to characterize the spectral, temporal and spatial specificity of the response modulations, as depicted in the main text figures.

### Cluster-based permutation statistics

We used cluster-based permutation statistics in the analysis of power relative to baseline as well as model coefficients. We determined cluster sizes as contiguous differences from zero with identical sign with p < 0.05 (random effects two-tailed t-test, uncorrected). The analysis was repeated 10,000 times, shuffling the signs of the effects per subject and taking the maximum cluster size determined as above. A cluster was determined to be significant at p < 0.05 when its size exceeded the 95th percentile of this maximum cluster size distribution to account for multiple testing (Nichols and Holmes 2002).

## Results

We recorded MEG in human subjects (n = 19) that viewed dynamic random-dot stimuli with varying luminance contrast (3 levels) and motion coherence (3 levels) (Fig. 1B). After stimulus presentation (1000 ms) and a variable 300 to 600 ms delay, the participants indicated whether they saw an upward or a downward motion with a button press (Fig. 1A).

In line with previous findings (Hall et al. 2005; Siegel et al. 2007; Hipp et al. 2011), stimulus presentation increased gamma-band activity (> 64 Hz) compared to the blank fixation baseline in the visual cortex contralateral to the visual stimulus (Fig. 2A; both *p* < 0.01, cluster-based permutation). For all further analyses we averaged responses across the sources within contralateral occipital cortex. The robust increase in the gamma band was accompanied by a more widespread decrease in lower frequency bands (Fig. 2A; all three frequency bands between 8 and 64 Hz *p* < 0.01, cluster-based permutation). Both, the high-frequency enhancement and low-frequency suppression started around 0.1 s post stimulus onset and were then sustained (Fig. 2B; both *p* < 0.01, cluster-based permutation).

**Figure 2.**
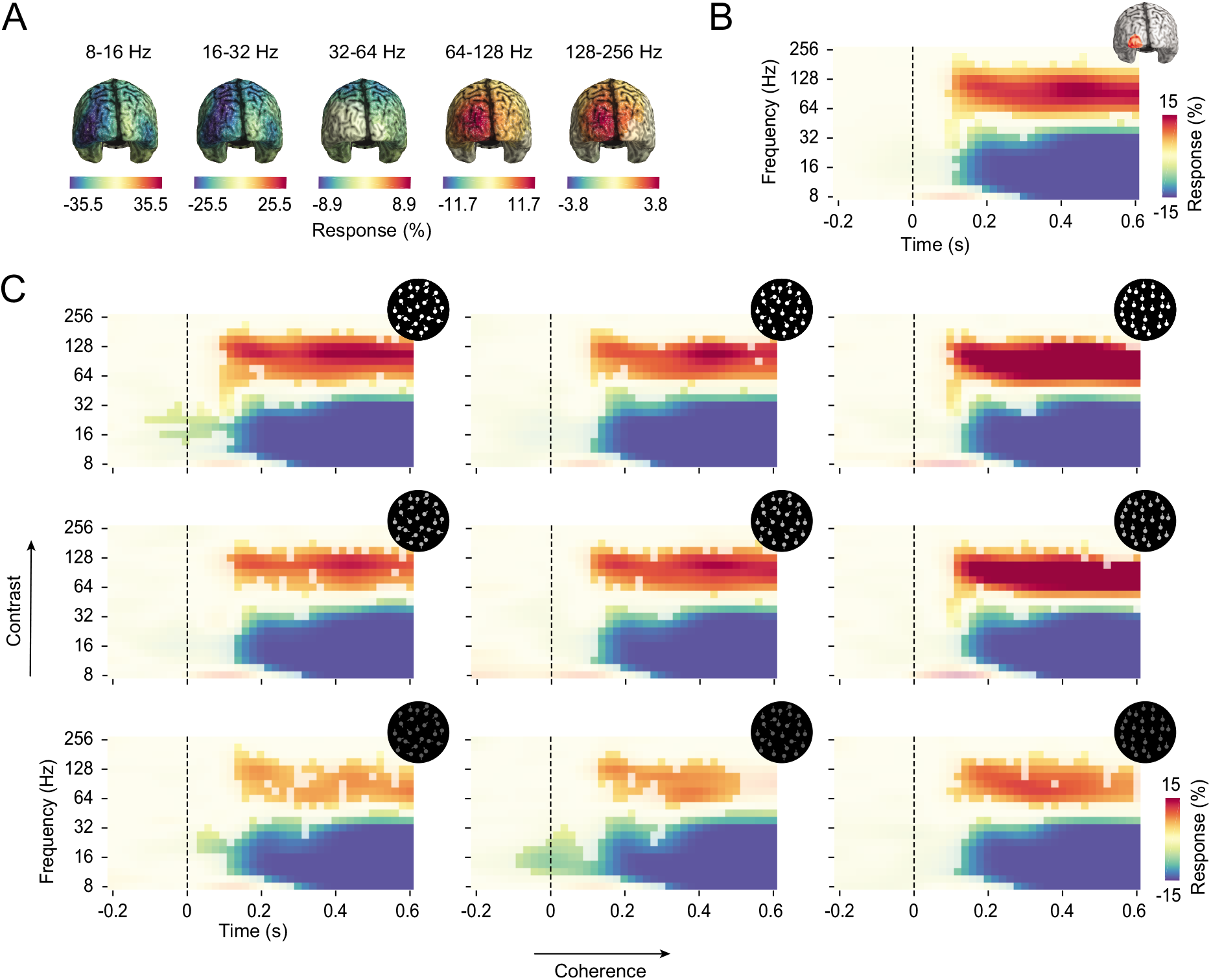
Stimulus induced responses relative to baseline. *A*, Cortical distribution of the power response during stimulation (0.1 to 1.1 s) in five frequency bands. Brains are viewed from the back. *B*, Power response resolved in time and frequency in occipital cortex contralateral to the visual hemifield of stimulation. The region of interest is depicted on the upper right. *C*, Time-frequency resolved power responses in each stimulus condition. In all panels, statistical significance (*p* < 0.05 corrected, cluster-permutation) is indicated by color opacity. Same color-scale as in *B*.

We next addressed how gamma-band activity varied with luminance contrast and motion coherence (Fig. 2C). For all 3 levels of motion coherence and contrast, visual stimulation induced a robust increase of gamma-band activity and decrease of low-frequency activity (Fig. 2C; all clusters *p* < 0.05, cluster-based permutation). Furthermore, gamma-band activity increased monotonically with both visual features (Fig. 2C). We next quantitatively assessed these modulations and tested for a potential interaction between coherence and contrast using a model-based approach.

We performed sequential polynomial regression to model the neuronal response as a linear and quadratic function of motion coherence and luminance contrast as well as of the interaction of these linear and quadratic features (Büchel et al. 1998; Rees et al. 2000; Siegel et al. 2007). The fitted model coefficients reveal the corresponding linear and quadratic modulations of the neuronal response by coherence and contrast (Fig. 3A). Importantly, stimulus coherence and contrast were uncorrelated by design, and all model coefficients were estimated independently using orthogonalized regressors. Thus, the interaction coefficients reflect multiplicative response modulations that cannot be explained by linear modulations (see Materials and Methods for further details). We found that both, contrast and motion coherence had a positive linear effect on visual gamma-band activity (Fig. 3A; contrast: *p* < 0.001; coherence: *p* < 0.0001, cluster-based permutation). These modulations were confined to frequencies from about 64 to 128 Hz, started shortly after stimulus onset and were then sustained throughout the stimulation period. In addition and in agreement with previous results (Gray and Singer 1989; Siegel and König 2003; Siegel et al. 2007), we observed a linear decrease of activity with contrast at lower frequencies from about 8 to 20 Hz (*p* < 0.01, cluster-based permutation). This effect started around 150 ms after stimulus onset. There was a weak sub-linear modulation (negative quadratic) of gamma-band activity with contrast (*p* < 0.05, cluster-based permutation). In addition and in line with previous results (Siegel et al. 2007), we observed a significant supra-linear (quadratic) increase of gamma-band activity with motion coherence between 32 and 128 Hz (*p* < 0.001, cluster-based permutation).

**Figure 3.**
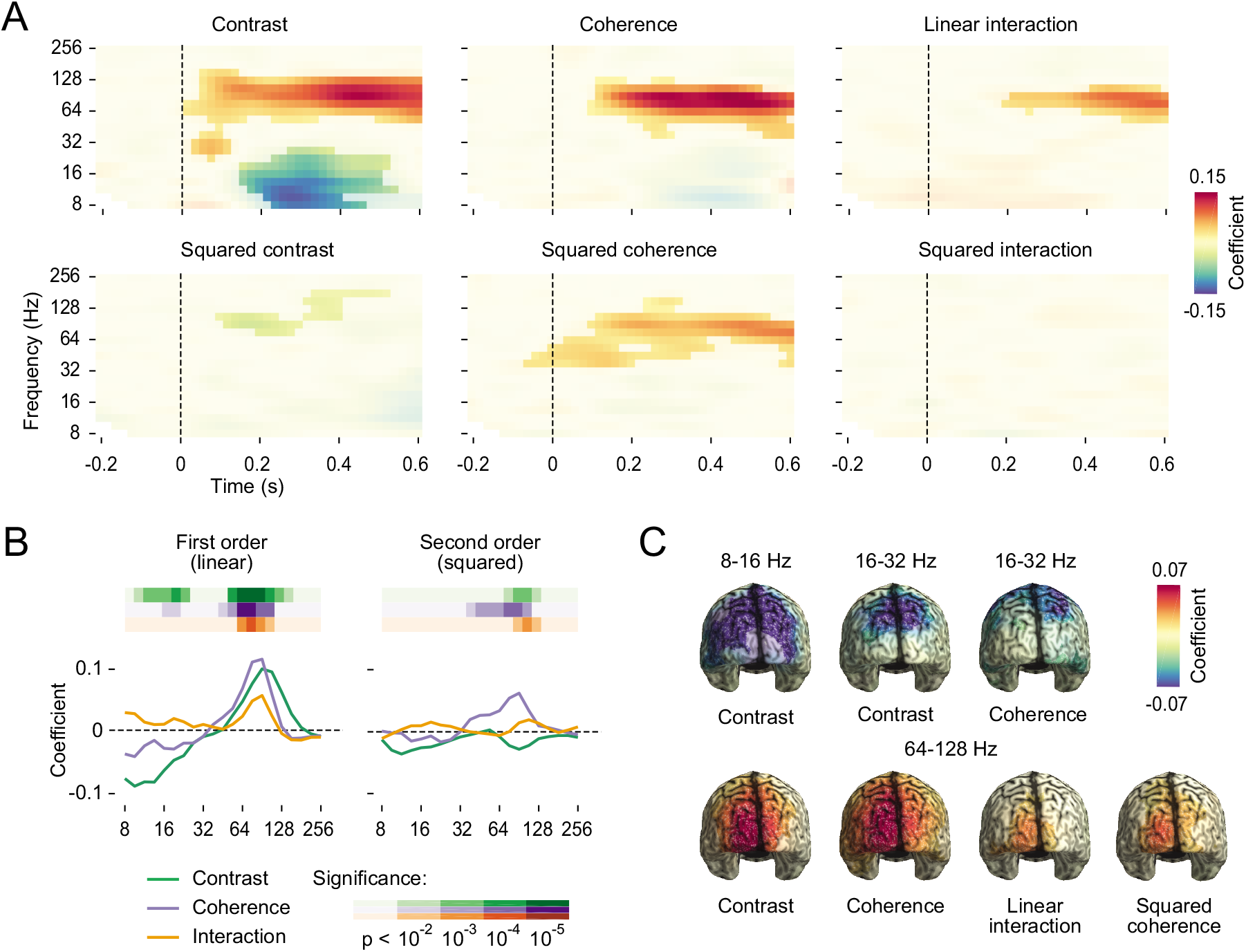
First- (linear) and second-order (quadratic) response modulations. *A*, Results of a sequential polynomial regression per time--frequency bin, including first- and second-order coefficients and their interactions. Statistical significance (*p* < 0.05 corrected, cluster-permutation) is indicated by color opacity. *B*, Response-coefficients as a function of frequency for the contralateral occipital cortex in the time window from 0.1 to 0.6 s past stimulus onset. Significant deviations from zero are indicated by the colored bars above the line-plots. *C*, Spatial specificity of selected response modulations in the time window from 0.1 to 0.6 s past stimulus onset. Statistical significance (*p* < 0.05 corrected, cluster-permutation) is indicated by color opacity. See Supplementary Fig. 1 for cortical distributions of all factors for all frequency bands.

Consistent with our hypothesis, we also found a robust multiplicative interaction between coherence and contrast (*p* < 0.01, cluster-based permutation, Fig. 3A). This interaction was confined to the frequency range between 64 and 128 Hz, started around 200 ms after stimulus onset and was then sustained. There was no such interaction between the negative response modulation of coherence and contrast at lower frequencies.

To further investigate the spectral profile of response modulations through coherence and contrast, we applied the same stepwise modelling (sequential polynomial regression) to the stimulus response averaged across time (0.1 to 0.6 s post stimulus onset) at each single frequency (Fig. 3B). In line with the temporally resolved analysis, this approach revealed robust linear effects of contrast and motion coherence in the gamma band and lower frequency ranges as well as quadratic effects of both visual features in the gamma-band (all *p* < 0.01, cluster-based permutation). We observed a multiplicative interaction between coherence and contrast in the gamma-band. Furthermore, in this analysis, we also observed a weak second-order interaction of contrast and coherence for the gamma-band (*p* < 0.01; multiplicative interaction of quadratic coherence and contrast). There was no significant linear or quadratic interaction of coherence and contrast for lower frequency ranges.

In which cortical regions do visual contrast and coherence modulate frequency specific neuronal population activity? To answer this question, we repeated the above analyses for each source location across the entire cortex, for 5 frequency bands (8-16 Hz, 16-32 Hz, 32-64 Hz, 64-128 Hz, 128-256 Hz) in the time range of 0.1 to 0.6 post stimulus and applied a spatial cluster-permutation statistic (Fig. 3C, 0.1 to 0.6 s post stimulus; all clusters shown *p* < 0.05, cluster-based permutation; see also Supplementary Fig. 1). Contrast and coherence induced a negative linear modulation in low frequencies (8 to 32 Hz) that extended along the dorsal visual stream peaking in occipitoparietal regions. The linear gamma-band modulations of contrast and coherence, including their multiplicative interaction, were more confined and shifted towards the pole of the occipital cortex (see Supplementary Fig. 1 for cortical distributions of all factors).

In a final step, we investigated how well the polynomial model fit the data. To this end, we repeated the regression analysis for neuronal responses in five frequency bands (Fig. 4; 0.1 to 0.6 s post stimulus; responses in contralateral visual cortex as above). Models of all but the middle frequency band (32 to 64 Hz) included contrast as a linear predictor (all *p* < 0.0001). Models between 16 and 128 Hz also included a linear coherence predictor (all *p* < 0.05). Both low frequency models (8 to 16 Hz and 16 to 32 Hz) included squared contrast (both *p* < 0.05), and for 32 to 64 Hz, the model included quadratic coherence (*p* < 0.05). Importantly, the linear interaction of coherence and contrast only improved the model for 64 to 128 Hz (*p* < 0.001). For this frequency range (64 to 128 Hz), the neuronal data was well fit (*R^2^* = 0.55) by the full response model including linear contrast, linear coherence, their interaction as well as quadratic contrast and coherence.

**Figure 4.**
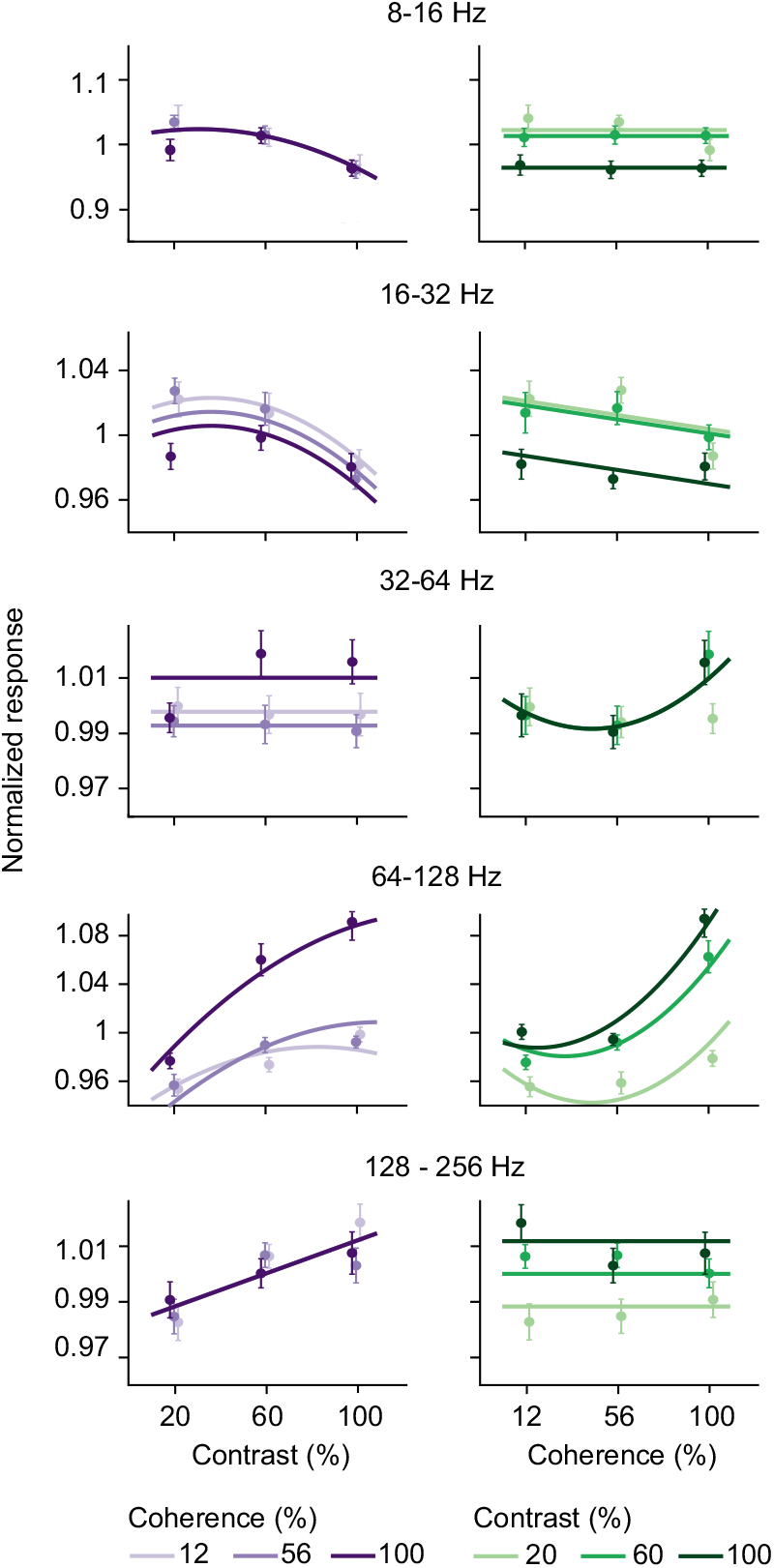
Average power response in the time window from 0.1 to 0.6 s in contralateral occipital cortex and fitted response models as a function of contrast, coherence and frequency band. Dots and error bars indicate the mean response and the corresponding standard error of the mean (SEM) across subjects. Lines indicate the fitted polynomial response model. Models include the significant coefficients (*p* < 0.05) of the sequential polynomial regression for the respective frequency band.

## Discussion

Here, we combined MEG, source reconstruction and parametric visual stimulation to test a critical prediction of the hypothesis that visual gamma-band activity acts as a cortical gain mechanism, which is that features that drive gamma-band activity interact super-linearly. To this end, we investigated the joined effect of visual contrast and motion coherence on gamma-band activity in human visual cortex. We found wide-spread activity modulations along the visual hierarchy in response to varying contrast and motion coherence. Low-frequency activity (8 to 32 Hz) decreased with coherence and contrast along the dorsal visual stream but exhibited no interaction between stimulus features. In contrast, for gamma-band activity, the driving influences of contrast and coherence interacted multiplicatively, thus confirming the prediction for a gain mechanism. Our findings provide novel evidence for the notion of gamma-band activity as a signature of local interactions that is driven through bottom-up sensory features and that regulates the gain or impact of sensory processing onto downstream regions (Siegel et al. 2007, 2008, 2012; Fries 2015).

The influence of contrast on neuronal spiking activity in visual cortex has been studied extensively. Neurons in areas along the dorsal visual stream exhibit a sigmoidal contrast response function (Albrecht and Hamilton 1982; Sclar et al. 1990; Martínez-Trujillo and Treue 2002), with different cells saturating at different contrast levels (Albrecht and Hamilton 1982). Gamma-band activity has been reported to increase approximately linearly with contrast in human MEG and EEG (Hall et al. 2005; Hadjipapas et al. 2015; Perry et al. 2015), while both linear (Logothetis et al. 2001; Henrie and Shapley 2005) and saturating (sub-linear) (Ray and Maunsell 2010a; Hadjipapas et al. 2015) modulations have been observed invasively in monkey visual cortex. In accordance with these reports, we found that gamma-band activity increased monotonically with contrast. Furthermore, we found that the increase of gamma-band activity with contrast was saturating (sub-linear), which accords well with recent results in non-human primates (Ray and Maunsell 2010b; Hadjipapas et al. 2015).

Previous studies observed a strong relationship between the peak-frequency of gamma-band activity and luminance contrast using grating stimuli (Ray and Maunsell 2010b; Hadjipapas et al. 2015). We did not observe such a frequency modulation in the present data (see Fig. 2). This may point to a stimulus specific origin of contrast-dependent frequency shifts in gamma-band activity (gratings vs. random-dot motion).

For motion coherence, response curves are also similar between single unit spiking and gamma-band population activity. The relationship between the motion coherence of a dynamic random-dot pattern and a cell’s response is predominantly linear (Britten et al. 1993; Heuer and Britten 2007). Also gamma-band activity in the human MEG increases approximately linearly with the motion coherence of dynamic random-dot patterns, with some subjects showing a quadratic (supra-linear) response (Siegel et al. 2007). Our results confirm these findings with both linear and quadratic modulations of gamma-band activity by motion coherence.

Although the relationship between single stimulus features and gamma-band activity has been studied extensively, little is known about the interaction of different stimulus features. Our results show that two stimulus features that monotonically increase gamma-band activity (contrast and motion coherence) interact supra-linearly in human visual cortex. This finding accords well with another MEG study that investigated the effect of three different stimulus parameters on gamma-band activity (full-field vs quadrant, static vs. motion, circular vs. linear grating) (Muthukumaraswamy and Singh 2013). Gamma activity exhibited main effects for all three stimulus features, and, in accordance with the present results, also significant positive interactions among all factors (Muthukumaraswamy and Singh 2013).

Local gamma-band activity likely arises from the interplay of both, lateral excitatory interactions and local inhibitory feedback (Bush and Sejnowski 1996; Kopell et al. 2000; Siegel et al. 2000; Bartos et al. 2007; Fries et al. 2007; Cardin et al. 2009; Fries 2009; Sohal et al. 2009; Donner and Siegel 2011; Vinck and Bosman 2016). The increase of gamma-band activity with contrast and motion coherence may reflect the enhanced rhythmic structuring of spiking activity with enhanced recruitment of these locally recurrent interactions through stronger bottom-up drive. Furthermore, stronger motion coherence enhances the spatiotemporal predictability of visual stimuli, which may further enhance the recruitment of stimulus specific lateral excitation (Gilbert and Wiesel 1989; Lund et al. 2003) and inhibition (Coen-Cagli et al. 2015; Vinck and Bosman 2016).

Visual gamma-band activity increases with selective visual attention (Fries 2001; Siegel et al. 2008), and enhances perceptual accuracy (Siegel et al. 2008) and response speed (Womelsdorf et al. 2006). Several factors may contribute to these behavioral effects. On the one hand, local gamma-band activity may rhythmically modulate and enhance the information content of neuronal spiking (Siegel et al. 2009; Womelsdorf et al. 2012; Vinck and Bosman 2016). On the other hand, local gamma-band activity may enhance the impact or gain of spiking activity on subsequent processing stages by to distinct mechanisms. First, the temporal synchronization of presynaptic spikes likely leads to their super-additive impact on postsynaptic neurons (MacLeod et al. 1998; Salinas and Sejnowski 2001; Azouz and Gray 2003; Laughlin and Sejnowski 2003; Fries 2009; Donner and Siegel 2011). Second, the rhythmic synchronization of presynaptic spiking may enhance its downstream impact by enabling its phase-alignment to corresponding postsynaptic rhythmic excitability fluctuations (Fries 2005; Siegel et al. 2008; Gregoriou et al. 2009; Bosman et al. 2012; Grothe et al. 2012). Our findings support this notion by showing a multiplicative interaction, i.e. an enhanced gain of gamma-band responses among visual features that drive this type of neuronal population activity.

Notably, we did not observe an interaction of coherence and contrast in their modulation of low-frequency activity (<30 Hz). Although, both features monotonically suppressed low-frequency activity in a graded fashion, there was no interaction between these effects. While ample evidence supports a behavioral effect of visual low-frequency population activity in particular in the alpha-band (Thut et al. 2006; Siegel et al. 2008; Jensen and Mazaheri 2010), our results suggest that, in contrast to gamma-band, slow rhythmic population activity may not exert a gain-like interaction between different visual features.

An interesting question is whether the different stimulus features modulated gamma-band activity preferentially in different cortical areas. Contrast may be expected to preferentially modulate earlier processing stages with steep contrast-response functions (e.g. V1), while motion coherence may preferentially modulate later stages specialized in motion processing, such as area MT+. Such differences may contribute to the multiplicative interaction in average gamma activity across visual cortex through a sequential gain enhancement across several processing stages. Although we observed modulations of gamma-band activity specifically in the contralateral visual cortex, due to the limited spatial resolution, it is difficult to pinpoint the exact cortical stages of gain modulation and interactions with MEG. Further invasive studies are required to address this question.

In sum, we find that visual motion and contrast interact multiplicatively in their drive of visual gamma-band activity. Gamma-band activity may reflect a visual gain mechanism that combines sensory features and regulates the impact of sensory processing onto downstream regions.

## Acknowledgements

This research was supported by the European Research Council (ERC) StG335880 (M.S), the Deutsche Forschungsgemeinschaft (DFG, German Research Foundation) project 276693517 (SFB 1233) (M.S.) and the Centre for Integrative Neuroscience (DFG, EXC 307) (M.S.).

## Author Contributions

Conceptualization: M.S.; Methodology: F.P., A.A.P., J.F.H., D.J.H., M.S.; Investigation: A.A.P., J.F.H.; Formal Analysis, F.P., D.J.H.; Writing – Original Draft, Review & Editing: F.P., D.J.H., M.S.; Funding Acquisition & Resources: M.S.; Supervision: M.S.

---

**Supplementary Figure 1.**
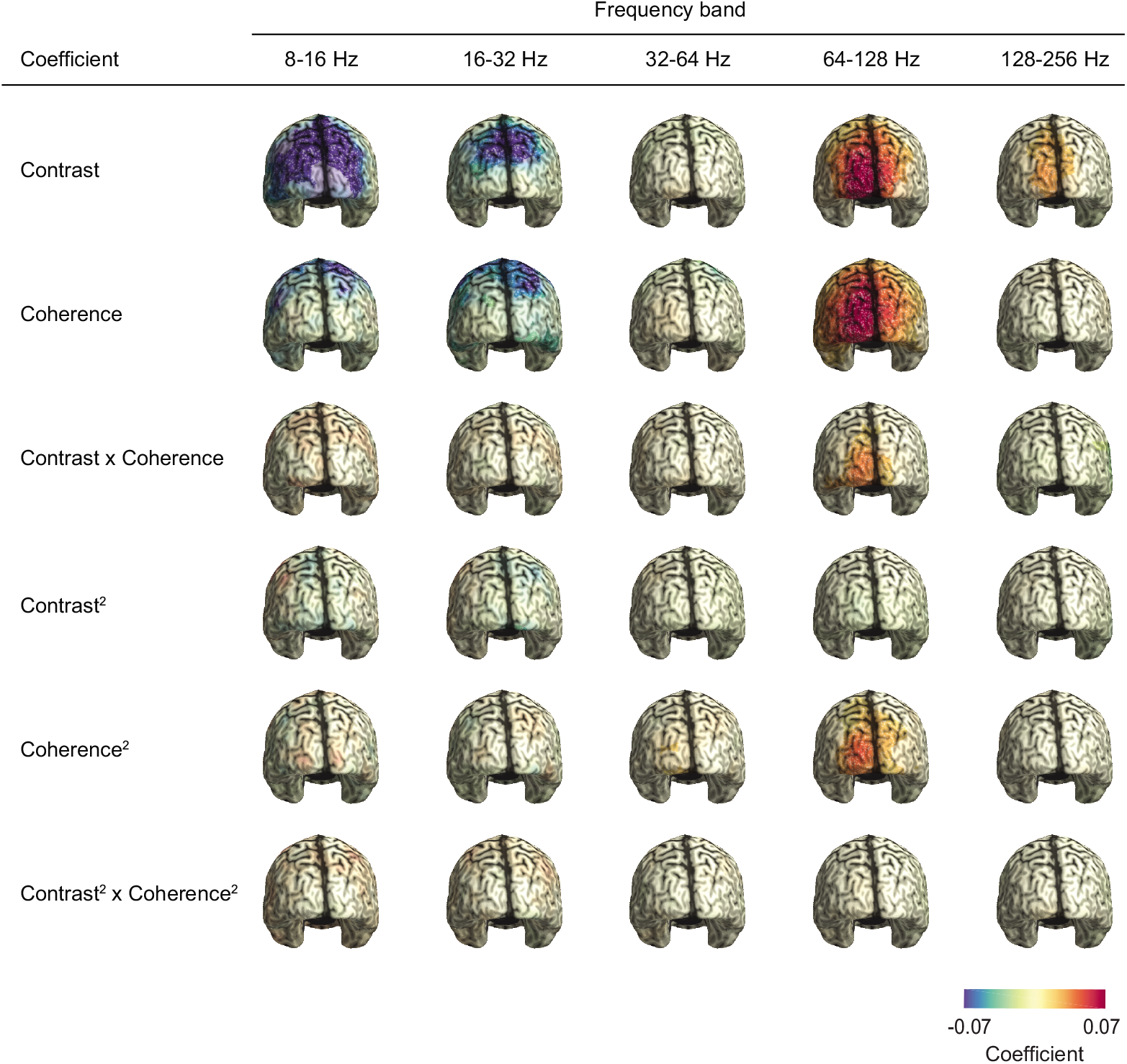
Cortical distribution of all response modulations in the time window from 0.1 to 0.6 s past stimulus onset for all five frequency-bands. Statistical significance (*p* < 0.05 corrected, cluster-permutation) is indicated by color opacity.

## References

Albrecht DG, Hamilton DB. 1982. Striate cortex of monkey and cat: contrast response function. Journal of Neurophysiology. 48:217–237.

Azouz R, Gray CM. 2003. Adaptive coincidence detection and dynamic gain control in visual cortical neurons in vivo. Neuron. 37:513–523.

Bartos M, Vida I, Jonas P. 2007. Synaptic mechanisms of synchronized gamma oscillations in inhibitory interneuron networks. Nat Rev Neurosci. 8:45–56.

Bosman CA, Schoffelen J-M, Brunet N, Oostenveld R, Bastos AM, Womelsdorf T, Rubehn B, Stieglitz T, De Weerd P, Fries P. 2012. Attentional stimulus selection through selective synchronization between monkey visual areas. Neuron. 75:875–888.

Britten KH, Shadlen MN, Newsome WT, Movshon JA. 1993. Responses of neurons in macaque MT to stochastic motion signals. Vis Neurosci. 10:1157–1169.

Büchel C, Holmes AP, Rees G, Friston KJ. 1998. Characterizing Stimulus–Response Functions Using Nonlinear Regressors in Parametric fMRI Experiments. NeuroImage. 8:140–148.

Bush P, Sejnowski T. 1996. Inhibition synchronizes sparsely connected cortical neurons within and between columns in realistic network models. J Comput Neurosci. 3:91–110.

Cardin JA, Carlen M, Meletis K, Knoblich U, Zhang F, Deisseroth K, Tsai LH, Moore CI. 2009. Driving fast-spiking cells induces gamma rhythm and controls sensory responses. Nature. 459:663–667.

Coen-Cagli R, Kohn A, Schwartz O. 2015. Flexible gating of contextual influences in natural vision. Nat Neurosci. 18:1648–1655.

Donner TH, Siegel M. 2011. A framework for local cortical oscillation patterns. Trends in Cognitive Sciences. 15:191–199.

Draper NR, Smith H. 1998. Applied regression analysis: includes disk. 3. ed., Wiley series in probability and statistics Texts and references section. New York, NY: Wiley

Friedman-Hill S. 2000. Dynamics of Striate Cortical Activity in the Alert Macaque: I. Incidence and Stimulus-dependence of Gamma-band Neuronal Oscillations. Cerebral Cortex. 10:1105–1116.

Fries P. 2001. Modulation of Oscillatory Neuronal Synchronization by Selective Visual Attention. Science. 291:1560–1563.

Fries P. 2005. A mechanism for cognitive dynamics: neuronal communication through neuronal coherence. Trends in Cognitive Sciences. 9:474–480.

Fries P. 2009. Neuronal Gamma-Band Synchronization as a Fundamental Process in Cortical Computation. Annual Review of Neuroscience. 32:209–224.

Fries P. 2015. Rhythms for Cognition: Communication through Coherence. Neuron. 88:220–235.

Fries P, Nikolić D, Singer W. 2007. The gamma cycle. Trends in Neurosciences. 30:309–316.

Gieselmann MA, Thiele A. 2008. Comparison of spatial integration and surround suppression characteristics in spiking activity and the local field potential in macaque V1. European Journal of Neuroscience. 28:447–459.

Gilbert CD, Wiesel TN. 1989. Columnar specificity of intrinsic horizontal and corticocortical connections in cat visual cortex. J Neurosci. 9:2432–2442.

Gray CM, Singer W. 1989. Stimulus-specific neuronal oscillations in orientation columns of cat visual cortex. Proceedings of the National Academy of Sciences. 86:1698–1702.

Gregoriou GG, Gotts SJ, Zhou H, Desimone R. 2009. High-frequency, long-range coupling between prefrontal and visual cortex during attention. Science. 324:1207–1210.

Gross J, Kujala J, Hamalainen M, Timmermann L, Schnitzler A, Salmelin R. 2001. Dynamic imaging of coherent sources: Studying neural interactions in the human brain. Proceedings of the National Academy of Sciences. 98:694–699.

Grothe I, Neitzel SD, Mandon S, Kreiter AK. 2012. Switching neuronal inputs by differential modulations of gamma-band phase-coherence. J Neurosci. 32:16172–16180.

Hadjipapas A, Lowet E, Roberts MJ, Peter A, De Weerd P. 2015. Parametric variation of gamma frequency and power with luminance contrast: A comparative study of human MEG and monkey LFP and spike responses. NeuroImage. 112:327–340.

Hall SD, Holliday IE, Hillebrand A, Singh KD, Furlong PL, Hadjipapas A, Barnes GR. 2005. The missing link: analogous human and primate cortical gamma oscillations. NeuroImage. 26:13–17.

Henrie JA, Shapley R. 2005. LFP Power Spectra in V1 Cortex: The Graded Effect of Stimulus Contrast. Journal of Neurophysiology. 94:479–490.

Heuer HW, Britten KH. 2007. Linear Responses to Stochastic Motion Signals in Area MST. Journal of Neurophysiology. 98:1115–1124.

Hipp JF, Engel AK, Siegel M. 2011. Oscillatory Synchronization in Large-Scale Cortical Networks Predicts Perception. Neuron. 69:387–396.

Hipp JF, Hawellek DJ, Corbetta M, Siegel M, Engel AK. 2012. Large-scale cortical correlation structure of spontaneous oscillatory activity. Nature Neuroscience. 15:884–890.

Hyvärinen A, Oja E. 2000. Independent component analysis: algorithms and applications. Neural Networks. 13:411–430.

Jensen O, Mazaheri A. 2010. Shaping functional architecture by oscillatory alpha activity: gating by inhibition. Front Hum Neurosci. 4:186.

Koelewijn L, Dumont JR, Muthukumaraswamy SD, Rich AN, Singh KD. 2011. Induced and evoked neural correlates of orientation selectivity in human visual cortex. NeuroImage. 54:2983–2993.

König P, Engel AK, Singer W. 1996. Integrator or coincidence detector? The role of the cortical neuron revisited. Trends Neurosci. 19:130–137.

Kopell N, Ermentrout GB, Whittington MA, Traub RD. 2000. Gamma rhythms and beta rhythms have different synchronization properties. Proc Natl Acad Sci U S A. 97:1867–1872.

Laughlin SB, Sejnowski TJ. 2003. Communication in neuronal networks. Science. 301:1870–1874.

Liu J, Newsome WT. 2006. Local Field Potential in Cortical Area MT: Stimulus Tuning and Behavioral Correlations. Journal of Neuroscience. 26:7779–7790.

Logothetis NK, Pauls J, Augath M, Trinath T, Oeltermann A. 2001. Neurophysiological investigation of the basis of the fMRI signal. Nature. 412:150–157.

Lund JS, Angelucci A, Bressloff PC. 2003. Anatomical substrates for functional columns in macaque monkey primary visual cortex. Cereb Cortex. 13:15–24.

MacLeod K, Bäcker A, Laurent G. 1998. Who reads temporal information contained across synchronized and oscillatory spike trains? Nature. 395:693–698.

Martínez-Trujillo JC, Treue S. 2002. Attentional Modulation Strength in Cortical Area MT Depends on Stimulus Contrast. Neuron. 35:365–370.

Muthukumaraswamy SD, Singh KD. 2013. Visual gamma oscillations: The effects of stimulus type, visual field coverage and stimulus motion on MEG and EEG recordings. NeuroImage. 69:223–230.

Nichols TE, Holmes AP. 2002. Nonparametric permutation tests for functional neuroimaging: A primer with examples. Human Brain Mapping. 15:1–25.

Niessing J. 2005. Hemodynamic Signals Correlate Tightly with Synchronized Gamma Oscillations. Science. 309:948–951.

Nolte G. 2003. The magnetic lead field theorem in the quasi-static approximation and its use for magnetoencephalography forward calculation in realistic volume conductors. Phys Med Biol. 48:3637–3652.

Oostenveld R, Fries P, Maris E, Schoffelen J-M. 2011. FieldTrip: Open Source Software for Advanced Analysis of MEG, EEG, and Invasive Electrophysiological Data. Computational Intelligence and Neuroscience. 2011:1–9.

Perry G, Hamandi K, Brindley LM, Muthukumaraswamy SD, Singh KD. 2013. The properties of induced gamma oscillations in human visual cortex show individual variability in their dependence on stimulus size. NeuroImage. 68:83–92.

Perry G, Randle JM, Koelewijn L, Routley BC, Singh KD. 2015. Linear Tuning of Gamma Amplitude and Frequency to Luminance Contrast: Evidence from a Continuous Mapping Paradigm. PLOS ONE. 10:e0124798.

Ray S, Maunsell JH. 2010a. Differences in gamma frequencies across visual cortex restrict their possible use in computation. Neuron. 67:885–896.

Ray S, Maunsell JHR. 2010b. Differences in Gamma Frequencies across Visual Cortex Restrict Their Possible Use in Computation. Neuron. 67:885–896.

Rees G, Friston K, Koch C. 2000. A direct quantitative relationship between the functional properties of human and macaque V5. Nature Neuroscience. 3:716–723.

Rohenkohl G, Bosman CA, Fries P. 2018. Gamma Synchronization between V1 and V4 Improves Behavioral Performance. Neuron. 100:953–963.e3.

Salinas E, Sejnowski TJ. 2001. Correlated neuronal activity and the flow of neural information. Nature Reviews Neuroscience. 2:539–550.

Scase MO, Braddick OJ, Raymond JE. 1996. What is noise for the motion system? Vision Res. 36:2579–2586.

Sclar G, Maunsell JH, Lennie P. 1990. Coding of image contrast in central visual pathways of the macaque monkey. Vision Res. 30:1–10.

Siegel M, Donner TH, Engel AK. 2012. Spectral fingerprints of large-scale neuronal interactions. Nature Reviews Neuroscience. 13:121–134.

Siegel M, Donner TH, Oostenveld R, Fries P, Engel AK. 2007. High-Frequency Activity in Human Visual Cortex Is Modulated by Visual Motion Strength. Cerebral Cortex. 17:732–741.

Siegel M, Donner TH, Oostenveld R, Fries P, Engel AK. 2008. Neuronal Synchronization along the Dorsal Visual Pathway Reflects the Focus of Spatial Attention. Neuron. 60:709–719.

Siegel M, König P. 2003. A Functional Gamma-Band Defined by Stimulus-Dependent Synchronization in Area 18 of Awake Behaving Cats. The Journal of Neuroscience. 23:4251–4260.

Siegel M, Kording KP, König P. 2000. Integrating top-down and bottom-up sensory processing by somato-dendritic interactions. J Comput Neurosci. 8:161–173.

Siegel M, Warden MR, Miller EK. 2009. Phase-dependent neuronal coding of objects in shortterm memory. Proc Natl Acad Sci U S A. 106:21341–21346.

Sohal VS, Zhang F, Yizhar O, Deisseroth K. 2009. Parvalbumin neurons and gamma rhythms enhance cortical circuit performance. Nature.

Thut G, Nietzel A, Brandt SA, Pascual-Leone A. 2006. Alpha-band electroencephalographic activity over occipital cortex indexes visuospatial attention bias and predicts visual target detection. J Neurosci. 26:9494–9502.

van Kerkoerle T, Self MW, Dagnino B, Gariel-Mathis M-A, Poort J, van der Togt C, Roelfsema PR. 2014. Alpha and gamma oscillations characterize feedback and feedforward processing in monkey visual cortex. Proc Natl Acad Sci USA. 111:14332–14341.

Van Veen BD, Van Drongelen W, Yuchtman M, Suzuki A. 1997. Localization of brain electrical activity via linearly constrained minimum variance spatial filtering. IEEE Transactions on Biomedical Engineering. 44:867–880.

Vinck M, Bosman CA. 2016. More Gamma More Predictions: Gamma-Synchronization as a Key Mechanism for Efficient Integration of Classical Receptive Field Inputs with Surround Predictions. Frontiers in Systems Neuroscience. 10.

Vinck M, Womelsdorf T, Fries P. 2013. Gamma-Band Synchronization and Information Transmission. In: Principles of Neural Coding. CRC Press. p. 449–470.

Womelsdorf T, Fries P, Mitra PP, Desimone R. 2006. Gamma-band synchronization in visual cortex predicts speed of change detection. Nature. 439:733–736.

Womelsdorf T, Lima B, Vinck M, Oostenveld R, Singer W, Neuenschwander S, Fries P. 2012. Orientation selectivity and noise correlation in awake monkey area V1 are modulated by the gamma cycle. Proc Natl Acad Sci USA. 109:4302–4307.

